# Cross-protection against Bundibugyo by Ebola and Sudan vaccines

**DOI:** 10.64898/2026.06.10.731377

**Authors:** Lara Kelchtermans, Viktor Lemmens, Johan Neyts, Yeranddy A. Alpizar, Kai Dallmeier

## Abstract

Ebola outbreaks continue to expand across Central Africa, yet available countermeasures remain limited and virus-specific. Whether available vaccines developed against the related Ebola virus (EBOV) and Sudan virus (SUDV) can confer cross-protection against Bundibugyo virus (BDBV) is incompletely understood. Here, we developed a BDBV surrogate challenge model and evaluated cross-protective immunity conferred by a set of monovalent vaccine candidates representing all three *Orthoebolavirus* species affecting humans. A yellow fever 17D-vectored BDBV vaccine (YF-BDB) expressing the BDBV glycoprotein (GP) as antigen conferred homologous protection in *Ifnar*^−^/^−^ mice, preventing systemic viral dissemination and providing survival following surrogate BDBV challenge. Intriguingly, also vaccination with YF-EBO or YF-SUD protected mice from such BDBV surrogate challenge. Conversely, while YF-BDB and YF-EBO fully protected against respective heterologous EBOV or BDBV infection, they conferred only partial protection against lethal challenge in a comparable SUDV model. Serological analyses revealed comparable titers of cross-reactive antibodies across all vaccine groups. However, neutralization, especially cross-neutralization, was limited suggesting a major role for non-neutralizing mechanisms in protection. These findings demonstrate asymmetry in cross-protective immunity among *Orthoebolavirus* vaccines and indicate that SUDV GP-based antigens may provide broader protection than EBOV- or BDBV-targeted candidates. However, our data add to the preliminary yet limited evidence that EBOV GP-based vaccines might offer at least some degree of protection against BDBV, the virus driving current Public Health Emergency of International Concern (PHIEC) in the Central-East Africa region.

## MAIN

Orthoebolaviruses cause recurrent outbreaks of severe disease across central Africa, with high case-fatality rates (40–90%) [1] and profound socioeconomic disruption [1,2]. Although epidemic preparedness has focused largely on the Ebola virus (EBOV), recent outbreaks caused by the Sudan virus (SUDV) and Bundibugyo virus (BDBV), that is currently re-emerging after 14 years of apparent dormancy, highlight the need for broader countermeasures [1].

With no licensed vaccine nor therapeutic available for BDBV, repurposing existing, readily available interventions against EBOV and SUDV represents a plausible scenario for containing the rapidly expanding 2026 epidemic in the Democratic Republic of the Congo and Uganda [3]. These include ERVEBO, a recombinant vesicular stomatitis virus (rVSV)-vectored vaccine expressing EBOV glycoprotein (GP) [4,5]; a similar rVSV-vectored vaccine expressing SUDV GP [6], Ad26-EBOV GP prime/MVA-vectored (ZABDENO/MVABEA) prime-boost regimens; and the monoclonal antibody therapies EBANGA and INMAZEB [7-9]. Unfortunately, preclinical and clinical data on cross-protection conferred by these interventions remain limited or non-conclusive; accordingly, the evidence base for their endorsement is weak.

Most licensed *Orthoebolavirus* vaccines or vaccines in preclinical and clinical development are monotypic, targeting a specific single virus species only [10]. Even previously licensed MVABEA booster, which encodes antigens from multiple filovirus species, primarily induces EBOV-specific immunity and has limited cross-species efficacy [7,11]. Similarly, licensed rVSV-EBOV vaccine ERVEBO fails to protect non-human primates (NHP) against SUDV [12,13], whereas partial cross-protection between EBOV and BDBV has been reported following either rVSV-EBOV vaccination or heterologous DNA/rAd5 immunization in NHP animal models [2,7,11]. In the latter setting, cross-reactive EBOV-induced CD8^+^ T cells, and some CD4^+^ T cells, yet no antibodies, have been implicated in protection [14]. Likewise, an experimental rVSV vaccine against Taï Forest virus, another member of the *Orthoebolavirus* genus, did not confer protection against BDBV despite inducing stronger BDBV-reactive antibody responses than rVSV-EBOV [12,13]. Consistent with marked antigenic differences between filovirus genera, virtually no cross-reactivity has been observed when analyzing immune responses to orthoebolaviruses with those seen in Marburg virus survivors or vaccinees [15,16].

Strategies to broaden filovirus coverage include ChAdOx-biEBOV, encoding both the EBOV glycoprotein (GP) and SUDV GP. However, protection against SUDV challenge in macaques remains incomplete [17]. Similarly, a blended vaccine, in which multiple monovalent vaccine candidates expressing different *Orthoebolavirus* antigens are combined in a single formulation, as exemplified with Ad26- and rVSV-vectored multivalent regimens [10,18,19] may result in immune interference and compromise the response to individual components [20].

Here, we evaluate the immunogenicity and efficacy of a previously established yellow fever 17D-vectored BDBV vaccine candidate (YF-BDB) [21] in a homologous surrogate challenge model. Using this toolbox, we further evaluate the extent to which monovalent vaccines expressing the GP of EBOV, SUDV, or BDBV confer cross-protection against infection and, potentially, fatal disease in a series of heterologous challenge models. These findings may help inform and rationalize antigen selection during current and future outbreaks, particularly as long as fully antigen-matched vaccines remain unavailable.

## RESULTS

### Development and Evaluation of a BDBV Challenge Model and Vaccine

Recombinant YF-BDB, expressing the BDBV GP from of live-attenuated YF17D vector (**Fig. 1a**), exhibited comparable growth kinetics to parental YF17D *in vitro* (**Fig. 1b**) and induced high titers of BDBV GP-specific IgG in hyperimmunized animals [21]. Leveraging on our previously established VSV-based surrogate models [21,22], we designed a VSV-based challenge virus expressing BDBV GP and a mCherry reporter (**Extended data Fig. 1a**). Rescue of fully infectious chimeric VSV-BDB virus was confirmed via mCherry expression and BDBV GP detection using serum from YF-BDB-vaccinated mice (**Extended data Fig. 1b**). *In vitro*, VSV-BDB replicated to lower titers than VSV-EBO (**Extended data Fig. 1c**) or previously described VSV-SUD [22]. Likewise, VSV-BDB was less pathogenic *in vivo*, seen that *Ifnar*^−^/^−^ mice animals challenged with 100 or 1000 PFU maintained body weight within 10% of baseline (**Extended data Fig. 1d-f**). Thus, VSV-BDB exhibited greater attenuation than previously established chimeric VSV viruses, as evidenced by a reduced potency to induce lethal disease.

**Figure 1.**
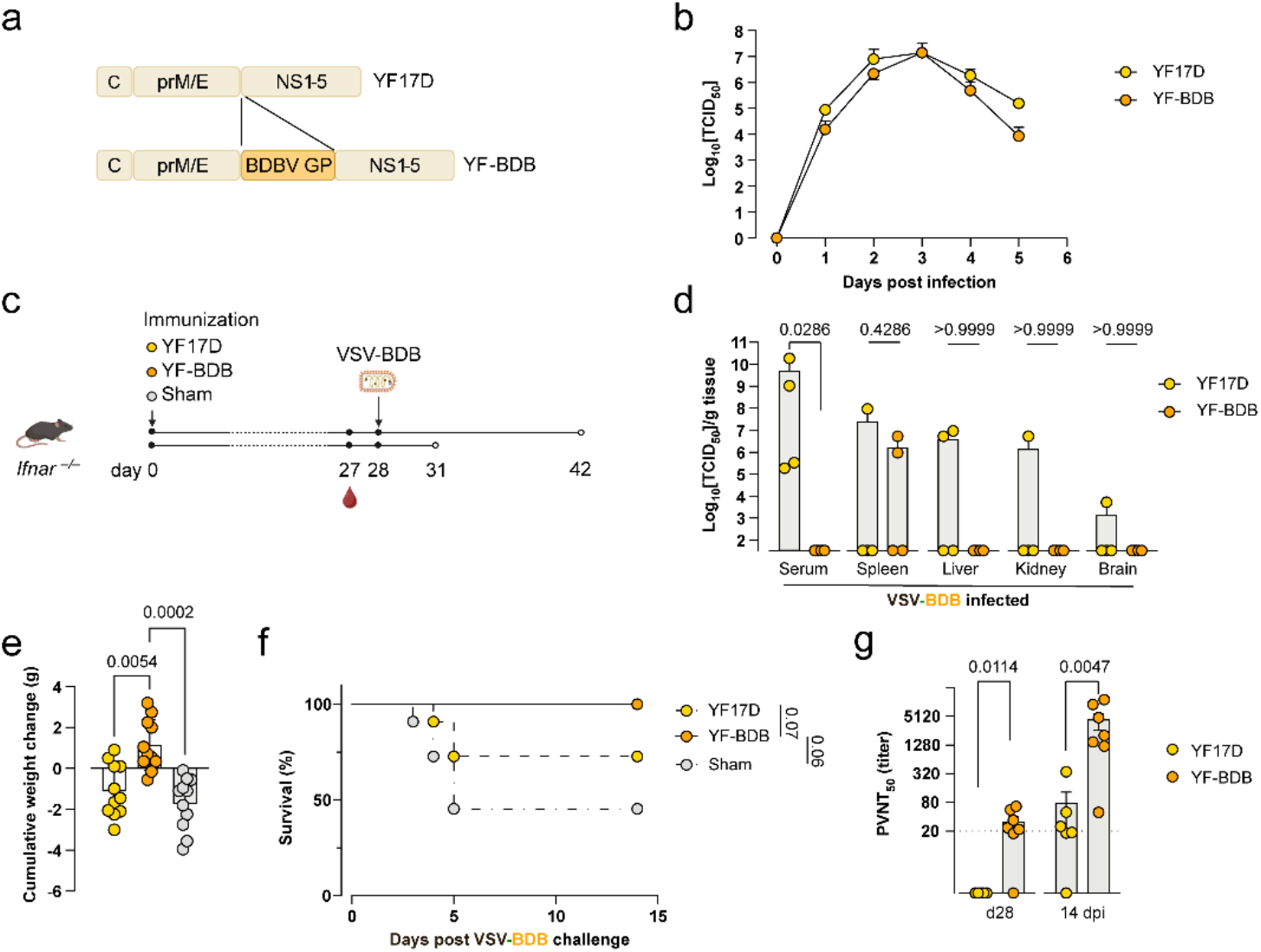
Development and immunogenicity of the YF-BDB vaccine candidate. **a**. Schematic representation of the YF-BDB vaccine construct. The Bundibugyo virus glycoprotein (BDBV GP) was inserted into the YF17D backbone at the junction site between the envelope (E) and non-structural protein 1 (NS1). **b**. *In vitro* replication kinetics of YF-BDB compared to parental YF17D in BHK21J cells (MOI = 0.01). Supernatants harvested until development of full cytopathic effect (CPE). Data represent mean titers ± SEM from two independent experiments. **c**. Schematic representation of the experimental design. *Ifnar*^−^/^−^ mice were vaccinated with YF17D (vector control), YF-BDB, or sham (2% medium). One month post-vaccination, mice were challenged with VSV-BDB. Animals were either monitored for survival until 14 days post-infection (dpi) or euthanized at 4 dpi for analysis of viral loads in specific tissues. **d**. Viral loads in target organs (serum, liver, kidney, brain, and spleen) of mice vaccinated with YF17D (n = 4) or YF-BDB (n = 4) at 4 dpi following VSV-BDB challenge. **e, f**. Cumulative weight change (**e**) and survival (Kaplan-Meier curve; **f**) of YF17D (n = 11), YF-BDB (n = 11), and sham (n = 11) vaccinated animals after VSV-BDB challenge. **g**. Neutralizing antibody titers against VSV-BDB. Data represent half-maximal pseudovirus neutralization titers (PVNT_50_) for YF17D and YF-BDB groups at day 28 post-vaccination and day 14 post-challenge (14 dpi). Data are presented as mean ± s.e.m. Statistical significance was determined using a two-sided Mann-Whitney U test (d, g), a two-sided Kruskal–Wallis test with Bonferroni correction (e), or a two-sided log-rank test for survival curves (f). *P*-values are indicated above the error bars.

Using this small animal model, we evaluated the efficacy of YF-BDB to confer homologous protection by vaccinating *Ifnar*^−^/^−^ mice with 250 PFU of YF-BDB, YF17D, or a sham control, followed by a VSV-BDB challenge 28 days later (**Fig. 1c**). By 4 days post-infection (dpi), YF-BDB-vaccinated mice had cleared the virus from the serum, liver, kidneys, and brain, with only 2 out of 4 mice showing transient virus presence in the spleen (**Fig. 1d**). In contrast, virus could readily be isolated at high titers from control animals, resulting in significantly higher disease scores at 4 dpi compared to the YF-BDB group (**Fig. 1e**). This finding further translated into 100% survival in YF-BDB-vaccinated mice, compared to only 60% and 80% survival in sham and YF17D controls, respectively (**Fig. 1f**). Moreover, YF-BDB elicited detectable neutralizing antibodies (nAbs) by 28 days post-vaccination, which were further boosted by 14 days post-challenge using a pseudovirus neutralization assay with replication competent VSV-BDB (**Fig. 1g**). These robust nAb responses likely contributed to, the rapid viral clearance observed in the tissues of YF-BDB-vaccinated animals.

### Cross-protection among BDBV, SUDV and EBOV can be achieved with monovalent vaccine candidates

To determine whether currently licensed vaccine for EBOV (ERVEBO) and trialed vaccine for SUDV (rVSV-SUDV) could elicit sufficient cross-reactive immunity to confer heterologous protection, we evaluated cross-protection amongst EBOV, SUDV and BDBV. To this end, *Ifnar*^−^/^−^ mice were immunized with previously described YF-EBO [22] and YF-SUD [22] vaccine candidates (2500 PFU) and infected 4 weeks post-vaccination with VSV-BDB (100 PFU). All vaccinated animals survived (**Fig. 2a**), showed minimal signs of disease **(Fig. 2b)** and a positive weight evolution **(Fig. 2c)** despite the heterologous challenge. By contrast, YF17D controls exhibited significantly higher disease scores and weight loss, which were virtually absent in YF-EBO and YF-SUD groups (*p*=0.0037and *p*=0.0214, respectively; **Fig. 2b-c, Extended data Fig. 2a**), and at least some, though low (10%) mortality. All in all, this confirms that both antigen-mismatched monovalent vaccines can provide cross-protection against virulent BDBV surrogate infection.

**Figure 2.**
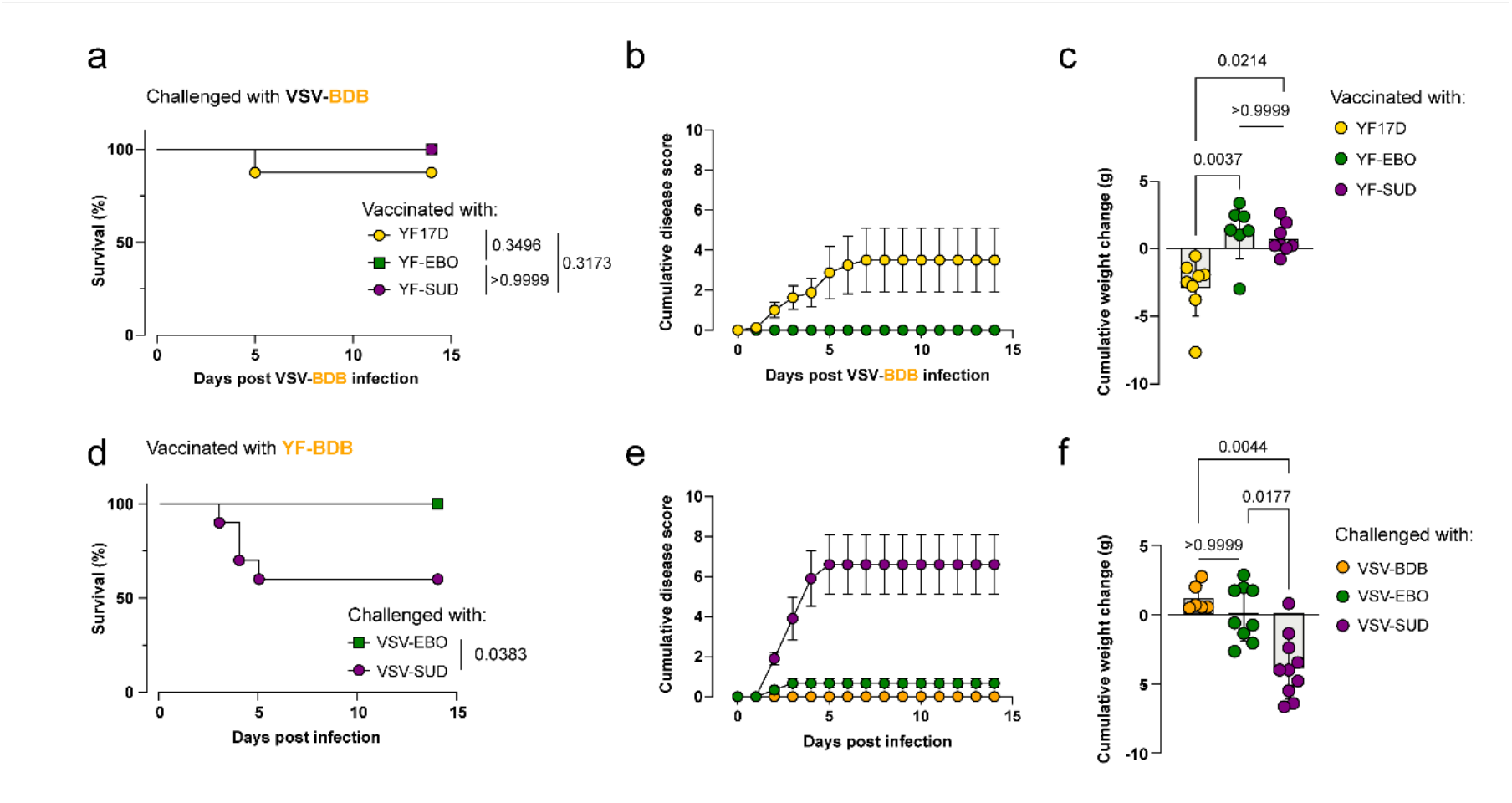
Cross-protection against BDBV (A-C) and heterologous efficacy of YF-BDB (D-F). **a-c**. Survival (Kaplan Meier curve; **a**), cumulative disease score (**b**), and cumulative weight loss of *Ifnar*^−^/^−^ mice vaccinated with YF17D (n= 8), YF-EBO (n= 7) and YF-SUD (n = 8) and challenged with VSV-BDB one month post-vaccination. **d-f**. Survival (Kaplan Meier curve; **d**), cumulative disease score (**e**), and cumulative weight change (**f**) of *Ifnar*^−^/^−^ mice vaccinated with YF-BDB and challenged with either VSV-BDB (n = 6), VSV-EBO (n = 9) or VSV-SUD (n = 10) one-month post vaccination. Data are presented as mean ± s.e.m. Statistical significance was determined using a two-sided Kruskal–Wallis test with Bonferroni correction (c, f), or a two-sided log-rank test for survival curves (a, d). *P*-values are indicated above the error bars.

To determine whether a BDBV GP vaccine could in turn confer cross-protection against EBOV or SUDV, we challenged YF-BDB-vaccinated animals with VSV-derived chimeric viruses expressing EBOV and SUDV GP. Indeed, vaccination using YF-BDB (2500 PFU) conferred full protection against lethal VSV-EBO infection (**Fig. 2d**) with no overt disease signs observed, except a transient weight loss of >5% in about half (5/9) of the mice (**Extended data Fig. 2b**), which resolved within 24 hours; translating accordingly into an only low disease score (**Fig. 2e**). Overall, YF-BDB-vaccinated animals appeared to cope similarly well with heterologous VSV-EBO infection as with VSV-BDB infection (e.g., *p*>0.9999 for cumulative weight change, **Fig. 2f**). Interestingly, YF-BDB achieved only 60% protection against VSV-SUD, in contrast to its high efficacy against VSV-EBO (**Fig. 2d**). YF-BDB-vaccinated mice challenged with VSV-SUD reached high disease scores (deferred grooming, hunched posture, vertebral segmentation, reduced mobility and limping in fatal cases; **Fig. 2e**) and a significant cumulative weight change (*p*=0.0044 compared to homologous VSV-BDB challenge, **Fig. 2f**) resulting in a weight loss of over 10% in 9 out of 10 animals (**Extended data Fig. 2b**). Altogether, these data suggest that a BDBV GP-based vaccine is unlikely to confer cross-protection against SUDV, while cross protection against EBOV is likely.

Previous findings are in line with the fact that the GP of EBOV is more similar to that of BDBV than SUDV GP (**Extended data Fig. 3a**,**b**). We therefore aimed to assess experimentally to which extent immunity between EBOV and SUDV overlaps. To that end, we vaccinated *Ifnar*^−^/^−^ mice with 2500 PFU of YF17D-based vaccine candidates expressing either EBOV GP (YF-EBO) or SUDV GP (YF-SUD) [21,22]. Four weeks post-vaccination, mice were challenged each with a lethal dose of either 100 PFU of VSV-EBO or VSV-SUD (**Fig. 3a**). In either case, 80-90% of YF17D-vaccinated animals included as controls succumbed within 3–5 dpi regardless of the challenge virus used (**Extended data Fig. 4a** *p=*0.2631), exhibiting high cumulative disease scores and significant weight loss within 3 dpi (**Extended data Fig. 4b-d**, *p=* 0.2065) As expected, YF-EBO conferred 100% protection against homologous VSV-EBO challenge [21]; however, it provided only 60% protection against heterologous VSV-SUD challenge (**Fig. 3b**, *p=* 0.2524), with both survivors and non-survivors showing elevated disease scores (**Fig. 3c**) and elevated cumulative weight changes (*p=* 0.0055, **Fig. 3d**), with weight loss exceeding 10% compared to their pre-infection weight (**Extended data Fig. 4e**).

**Figure 3.**
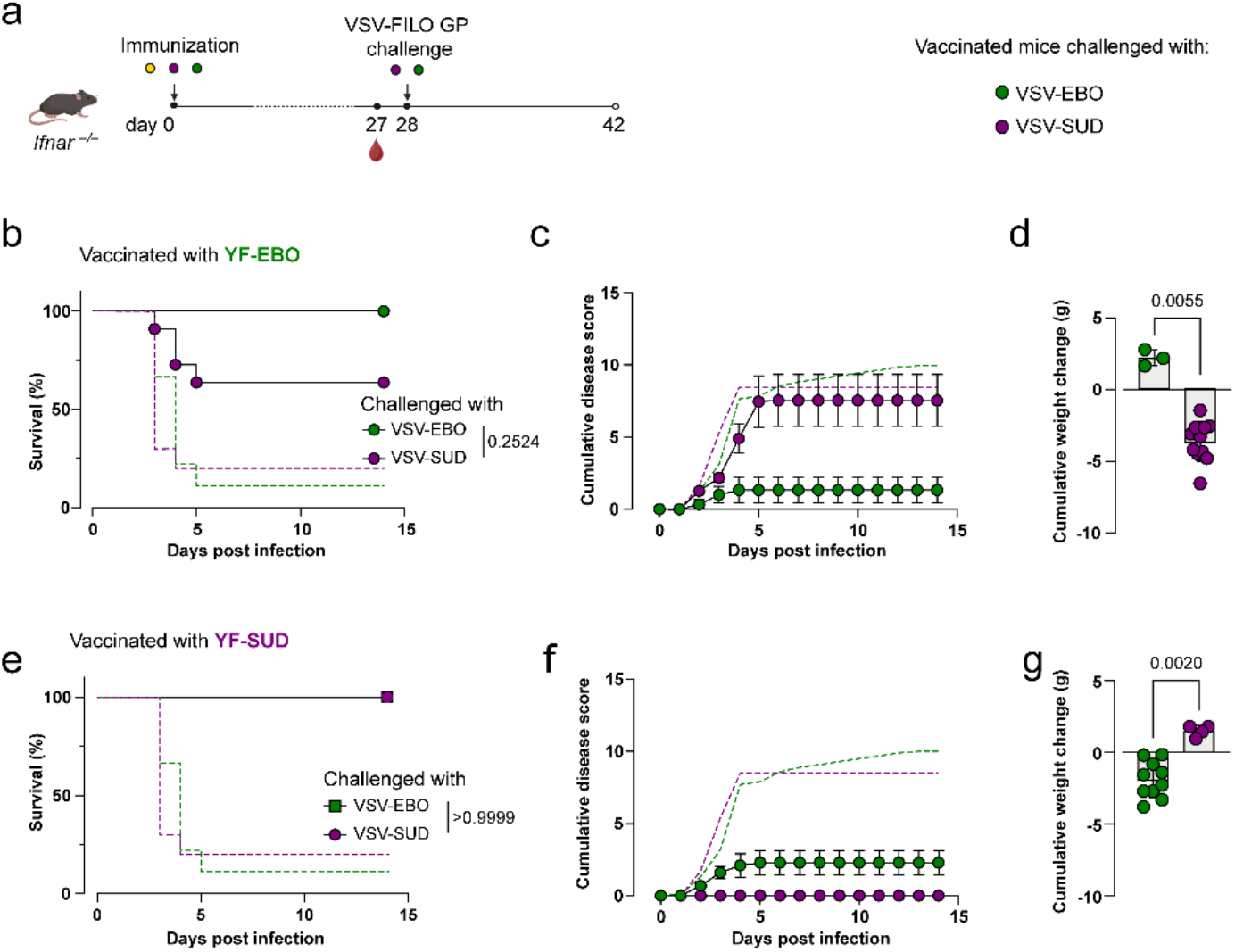
YF-SUD confers superior cross-protection against heterologous VSV-EBO challenge than YF-EBO to VSV-SUD. **a**. Schematic representation of the experimental design: *Ifnar*^−^/^−^ mice were vaccinated with YF17D (vector control, yellow), YF-SUD (purple), or YF-EBO (green). One month post-vaccination, mice were challenged with 100 PFU of either VSV-SUD or VSV-EBO. Clinical signs and weight changes were monitored until experimental endpoint at 14 days post-infection. **b-d**. Survival (**b**), cumulative disease score (**c**), and cumulative weight change of YF-EBO-vaccinated mice, following challenge with VSV-EBO (green datapoints with full line, n = 3) or VSV-SUD (purple datapoints with full line, n = 10). Survival and cumulative disease score data from the YF17D vector control is represent by the dotted line (green, VSV-EBO; purple, VSV-SUD). **e-g**. Survival (**e**), cumulative disease score (**f**), and cumulative weight change (**g**) of YF-SUD-vaccinated mice, following challenge with VSV_-_EBO (green datapoints with full line, n = 10) or VSV-SUD (purple datapoints with full line, n = 4). Survival and cumulative disease score data from the YF17D vector control is represent by the dotted line (green, VSV-EBO; purple, VSV-SUD). Data are presented as mean ± s.e.m. Statistical significance was determined using a two-sided Mann-Whitney U test for cumulative weight changes (d, g), or a two-sided log-rank test for survival curves (b, f). *P*-values are indicated above the error bars.

In contrast, YF-SUD vaccination elicited complete protection (100% survival) against both homologous (VSV-SUD) as well as heterologous (VSV-EBO) infection (**Fig. 3e**, *p>* 0.9999). Although YF-SUD-vaccinated animals challenged with VSV-EBO had a slightly higher cumulative disease score, driven by body weight changes, no overt clinical signs were observed regarding physical appearance or behavior (**Fig. 3f**). Moreover, these animals showed a significantly greater cumulative weight change than those receiving a homologous challenge (**Fig. 3g**, *p=*0.0020) while maximal weight loss remained limited to 10% compared to pre-infection values (**Extended data Fig. 4f**).

### Neutralizing Antibodies play a negligible role in protection

We next determined virus-specific IgG antibody titers, as antibodies specific for respective filovirus GP are known to correlate with protection [23]. All three vaccines (YF-EBO, YF-SUD, and YF-BDB) induced comparable titers of antibodies reacting with EBOV GP and BDBV GP (**Fig. 4a**). In contrast, YF-EBO failed to elicit detectable seroconversion to heterologous SUDV GP, and YF-BDB induced only very low SUDV GP-specific titers. Only YF-SUD elicited robust binding antibodies to SUDV GP, its homologous antigen (**Fig. 4a**).

**Figure 4.**
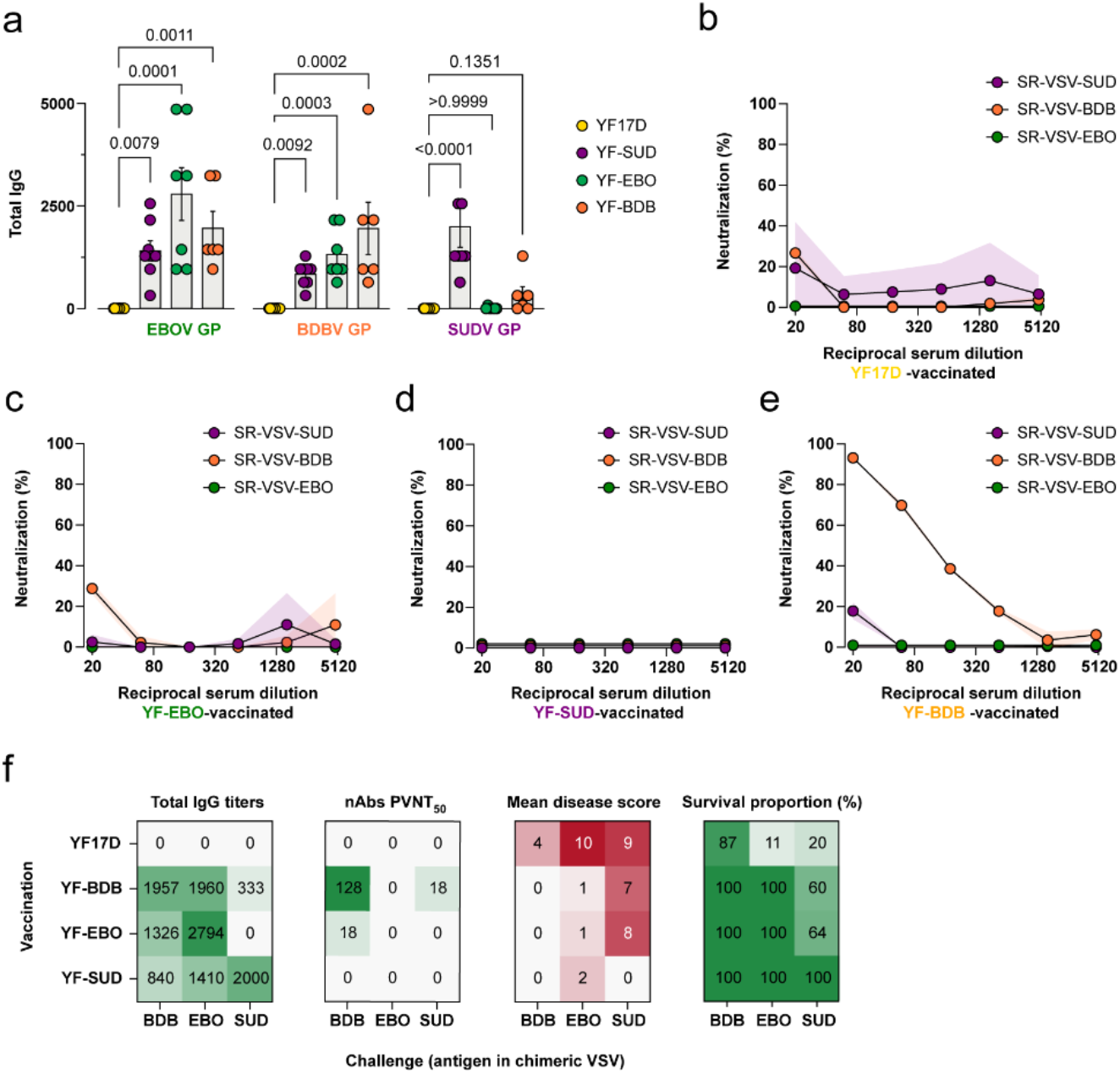
Cross-reactive antibodies do not translate into cross-neutralization. **a**. (Cross-)reactive binding IgG titers from sera of YF17D (n = 10), YF-SUD (n = 8), YF-EBO (n = 7), and YF-BDB (n = 6) vaccinated animals measured 28 days post-vaccination specific for EBOV GP, SUDV GP, and BDBV GP. **b-e**. Neutralizing activity (%) plotted against reciprocal serum dilutions for YF-EBO (**b**), YF-SUD (**c**), YF17D (**d**), and YF-BDB (**e**) hyperimmune sera. Neutralization was assessed using single-round VSV (SR-VSV) particles expressing either SUDV GP, BDBV GP, or EBOV GP. Data represent mean ± s.e.m. from three independent replicates; error bars are indicated by the colored shading of each line. **f**. Heatmaps representing cross-reactivity observed among EBOV, SUDV, and BDBV for binding antibody titers, PVNT_50_ neutralization titers, disease score and survival. Data are presented as mean ± s.e.m. Statistical significance was determined using a two-sided Kruskal–Wallis test with Bonferroni correction for multiple comparisons. *P*-values are indicated above the error bars.

Previously, YF-EBO and YF-SUD were used at a low dose of once 250 PFU yet failed to elicit detectable neutralizing antibodies (nAbs) of any relevance for immunity [21,22]. For the cross-protection experiments we used a 10-fold higher dose, i.e. 2500 PFU. To determine whether this could finally induce nAbs, a pseudovirus neutralization assay using single-round infectious VSV (SR-VSV) particles was established. Furthermore, to test whether the absence of detectable nAbs was related to the still relatively low, single-dose vaccination regimen used throughout our efficacy studies [21,22], a hyperimmunization regimen (three doses of 5,000 PFU) was applied. Even under these conditions, YF-EBO and YF-SUD failed to induce nAbs against EBOV, SUDV, or BDBV SR-VSV particles (**Fig. 4b**,**c**), similar to baseline YF17D vector control (**Fig. 4d**). In contrast, YF-BDB neutralized only SR-VSV-BDB GP, but did not cross-neutralize EBOV or SUDV GP SR-VSV particles (**Fig. 4e**). Altogether, these data suggest that nAbs contribute minimally to (cross-)protection across different orthoebolaviruses in this model.

Altogether, vaccine based on the GPs of BDBV and EBOV are unlikely to confer cross-protection against SUDV. However, it is suggestive that both licensed EBOV and SUDV vaccines could at least partially protect against BDBV; with implications for the ongoing Bundibugyo epidemic. Even more, SUDV-targeting vaccines may provide broad protection against a wider range of, if not all, orthoebolaviruses.

## DISCUSSION

In this study, we characterized a YF17D-vectored BDBV vaccine candidate and demonstrated that cross-protection to BDBV is achievable with EBOV or SUDV-targeting vaccines. Latter finding could be of particular interest amidst the ongoing BDBV outbreak in Central-East Africa of unprecedented magnitude, with no vaccines nor therapeutics available for (prophylactic) treatment. With the disclaimer that our studies were conducted using chimeric VSV to mimic genuine *Orthoebolavirus* infections, our data suggests an asymmetry in cross-protection among *Orthoebolavirus* vaccine candidates (**Fig. 4f**). While the SUDV GP-targeted vaccine (YF-SUD) conferred complete protection against a heterologous EBOV- and BDBV-like challenge, the reciprocal protection provided by EBOV (YF-EBO) and BDBV (YF-BDB) vaccines against the SUDV surrogate was significantly limited. This hierarchy of protection suggests that the SUDV GP may elicit a broader and more resilient immune profile than its EBOV and BDBV counterparts, providing a potential template for a “pan-*Orthoebolavirus*” vaccine strategy. Moreover, our data also suggest, similar to what has been observed for ERVEBO (rVSV-EBOV) in NHPs [15,24], that EBOV GP vaccine candidates might confer at least partial protection against BDBV.

The limited cross-reactivity of YF-EBO-induced antibodies toward SUDV is consistent with the structural landscape of the ebolavirus glycoprotein. Antibodies can access and bind comparable epitopes on BDBV and EBOV [25], despite the exact binding footprint differing among species [20,26,27]. EBOV GP-induced responses predominantly target the highly divergent Mucin-Like Domain (MLD), eliciting antibodies that are typically non-neutralizing [28,29]. Whereas more conserved regions such as the fusion loop and GP base, may offer targets for broader cross-reactive immunity [30,31]. YF-BDB was uniquely capable of eliciting homologous nAbs. This is likely supported by the greater structural flexibility and conformational heterogeneity of the BDBV GP, which renders neutralizing epitopes at the GP base more accessible than in the more rigid EBOV and SUDV glycoproteins [26]. This aligns with clinical data showing that BDBV survivors develop potent neutralizing responses targeting the GP base rather than the MLD decoy [32]. While our BDBV vaccine candidate successfully induced these homologous nAbs, they lacked cross-neutralizing breadth. This is likely explained by specific biochemical barriers at key neutralizing sites; for example, the fusion-loop pocket of EBOV GP possesses a more basic electrostatic charge than that of BDBV, preventing BDBV GP-specific antibodies from binding heterologous species [33].

We constructed three rVSV-based surrogate viruses expressing the glycoproteins of EBOV, SUDV, or BDBV. While the high mortality rates for the EBOV and SUDV constructs in the *Ifnar*^−^/^−^ model were comparable to those that can be observed for the original viruses in human outbreaks [30], the VSV-BDB virus failed to induce severe disease. Interestingly, increasing the infectious dose from 100 PFU to 1000 PFU did not increase mortality. Conversely, the animals exhibited a reduced morbidity at the higher dose. One hypothesis is that the high-dose inoculum may facilitate the rapid generation of defective interfering (DI) particles, which are known to dampen rVSV replication and promote a self-limiting, abortive infection [34], possibly involving the induction of an IFNα/β-independent antiviral state [35]. Nevertheless, the fact that VSV-BDB exhibits a significantly lower pathogenicity compared to EBOV and SUDV constructs sharing the same rVSV backbone, despite similar viral loads in organs at peak virus infection, further underscores that GP may serve as a primary driver of filovirus virulence [36,37].

The 70% sequence variability between EBOV and SUDV GP (**Extended data Fig. 3a**,**b**) has traditionally been viewed as a major hurdle for cross-protection. However, our results challenge the notion that sequence identity alone can be taken as reliable predictor of vaccine efficacy. In line, previous studies noted that a Taï Forest virus GP-based vaccine candidate failed to confer cross-protection against BDBV, despite their close genetic relatedness [15]. Similarly, we demonstrate that notwithstanding the significant antigenic distance, our SUDV vaccine candidate effectively bridged the gap to both EBOV and BDBV surrogate challenges, maybe by any combination of its multiple molecular and cellular mechanisms. Yet, also our EBOV vaccine candidate conferred equally cross-protection against BDBV.

### Limitations of the study

A major limitation of this study is the reliance on rVSV surrogate viruses expressing *Orthoebolavirus* GPs for the challenge experiments. While rVSV-GPs are invaluable tools for evaluating GP-targeted vaccine candidates in BSL-2 settings, they do not fully recapitulate the virus morphology, complex pathogenesis, systemic disease progression, or multisystem organ failure characteristic of original *Orthoebolavirus* infections. However, because our YF17D-vectored candidates exclusively target the GP, which is natively presented on the surface of rVSV particles, this model provides a highly relevant system for assessing GP-specific humoral protective mechanisms in a preclinical small-animal model setting. More importantly, our surrogate model is based on a robust challenge virus replication and spread yielding a multi-organ pathology. Therefore, even in the non-lethal BDBV GP model, we see a clear difference in phenotype between vaccinated and sham-vaccinated animals. Moreover, despite the stringency of the here employed challenge conditions, we can show protection among the different viruses.

## Conclusion

Previous clinical efforts have primarily focused on EBOV-centric vaccines, largely driven by the scale of the 2014–2016 West African outbreak. In contrast, SUDV and BUDV have received comparatively little attention until their recent re-emergence in 2025 [6] and 2026 [38]. Although not standard practice, the repurposing of existing vaccines may be considered in the absence of virus-specific countermeasures. In this context, EBOV GP-based vaccines could potentially confer cross-protection against BDBV. However, our findings suggest that a SUDV-based vaccine candidate might offer a geographically and virologically more versatile tool for preparedness and outbreak control. Future studies should focus on identifying and characterizing specific broadly cross-reactive vaccine targets such as conserved T cell epitopes and the non-neutralizing antibody landscape that appear to facilitate such robust cross-protection.

## METHODS

### Cells

BHK21J, Vero E6 cells, and VeroE6-miRFP670, were cultured in seeding medium, containing Minimal Essential Medium (MEM, Gibco) supplemented with 10% fetal calf serum (FCS, Gibco, 1% NEAA (Gibco), 1% L-Glutamine (Gibco), 1% NaHCO_3_ (Gibco), and 1% penicillin-streptomycin (Pen-Strep, Gibco). HEK293TAd cells were cultured in Dulbecco’s Modified Eagles Medium (DMEM, Gibco) seeding medium supplemented with 10% FCS (Gibco), 1% L-Glutamine (Gibco), 1% NaHCO_3_ (Gibco), 1% Pen-Strep (Gibco). Assay medium consisted of seeding medium with reduced concentration of 2% FCS.

### Constructs

YF-SUD, YF-EBO and YF-BDB were designed as previously described [21,39], by inserting the respective filovirus GP into the backbone of YF17D. Replication competent VSV-SUD, VSV-EBO and VSV-BDB was generated by inserting the respective GP, VSV intergenic sequence and fluorescent protein into pVSV-ΔG-GFP-2.6 (Extended data Fig. 1a) [21,22,40], kindly provided by Prof. Dr. M. A. Whitt, using *Mlu*I and *Avr*II restriction sites. The chimeric viruses were rescued using the methodology as previously described [40].

Single-round infectious VSV (SR-VSV) pseudotypes were generated by seeding 7.5×10^6^ HEK293T cells in a T75 flask. The following day, the culture medium was replaced with assay medium, and cells were transfected with 25 µg of pCMV-EBOV-GP-IRES-RFP/ pCMV-BDBV-GP-IRES-RFP/ pCMV-SUDV-GP-IRES-RFP plasmid using TransIT™-LT1 (Mirus Bio) according to the manufacturer’s instructions. After 24 h of incubation (37 °C, 5 % CO_2_), transfected cells were infected with ΔG-VSV* complemented with VSV-G (Indiana serotype) at a MOI 3. Following a 2 h absorption period (37 °C, 5 % CO_2_), the inoculum was removed and replaced with seeding medium containing a 1:250 dilution of L1-hybridoma anti–VSV-G antibody to neutralize residual input virus. After an additional 24 h of incubation (37 °C, 5 % CO_2_), the supernatant containing SR-VSV pseudotypes was collected and clarified by centrifugation at 15,000 rpm for 5 min.

### Vaccine virus stocks

Yellow fever 17D virus stock was obtained by passaging YF17D (Stamaril®, Sanofi-Pasteur MSD) twice in T75 flasks with 4.5 × 10^6^ Vero E6 cells. Virus stocks were harvested at 3 days post-infection after full cytopathogenic effect (CPE). YF-SUD, YF-BDB, and YF-EBO were obtained by transfecting 20 µg of pShuttle-YF17D-orthoebolavirus GP (either SUDV GP, BDBV GP or EBOV GP) into BHK21J cells (3.5 × 10^6^) seeded the day before in a T75 flask, using TransIT-LT1 Transfection Reagent (Mirus Bio, Merck), according to the manufacturer’s protocol. Prior to transfection, the seeding medium was exchanged for assay medium, and the vaccine virus was harvested at 4 days post-transfection, when full CPE was observed.

### Growth curves

BHK21J cells (3 × 10^6^ cells) for vaccine viruses, and Vero E6 cells (3 × 10^6^ cells) for VSV-viruses were seeded in T25 flasks to assess the growth curves of the YF-BDB, VSV-EBO, and VSV-SUD virus, respectively. After 24 h, the medium was replaced with assay medium and the cells were infected with 3 × 10^4^ PFU of YF-SUD, YF17D, VSV-BDB, or VSV-EBO. The virus was incubated with cells for 1 h at room temperature (RT), followed by thorough washing with seeding medium. Virus-containing medium (1 mL) was collected daily, with each collection replaced by 1 mL fresh assay medium, until full CPE was observed.

### Animals

*Ifnar*_*–/–*_ (B6.129S2-Ifnar1^tm1Agt^) mice of mixed gender were housed and bred in individually ventilated cages (max 5/cage) under SPF conditions with a 12/12 light/dark cycle (room temperature: 22 °C ± 2; humidity: 45-70%). Animals received food and water *ad libitum*, and cage enrichment to encourage normal behavior. Bedding material was weekly renewed, and all materials were autoclaved. Animal experiments were approved by the KU Leuven Ethical Committee (project numbers P140/2016, p100/2019, p164/2022, and p137/2024) and reported herein in accordance with ARRIVE (Animal Research: Reporting of In Vivo Experiments) guidelines. Experimental groups were mixed within cages to reduce cage effects. Sample sizes were defined *a priori*, by performing a power analysis with the expected effect size, statistical power of 80% and a significance level of 0.05, using G power and Sample size calculator Version 1.062 of the Universität Wien [41-43]. Assignment of the groups were blinded for operators, except for the lead investigator. For all experiments, the body weight and health status of the animals were daily monitored, and the clinical score was assessed according to Institutional Animal Care and Use Committee (IAUCUC) guidelines [44], and euthanized at humane endpoint. Cumulative weight loss was calculated for each individual animal by determining the area under the curve of weight change (g) during the first 3 (VSV-EBO, VSV-SUD) or 4 (VSV-BDB) days after challenge. Mice with abnormal development were excluded after weaning.

#### In vivo characterization of VSV-BDB

*Ifnar*^−^/^−^ mice (10-12-week-old) were infected by intraperitoneal (i.p.) administration of 100 PFU or 1000 PFU of VSV-BDB. Survival and body weights were monitored for the following 6 days.

#### Survival studies

*Ifnar*^*–/–*^ mice or (6–8-week-old) were vaccinated with 2500 PFU of YF17D, YF-SUD, YF-EBO, or YF-BDB. Mice were challenged i.p. at 1 month post-vaccination with 100 PFU VSV-SUD, VSV-EBO, or VSV-BDB. Blood was drawn submandibularly 2 days prior to challenge. After infection, water bottles were fitted with an extended nipple to facilitate access to water.

#### Viral loads

*Ifnar*^*–/–*^ mice or (6–8-week-old) were vaccinated with 2500 PFU of YF17D or YF-BDB and were challenged at 1 month post-vaccination with either VSV-BDB or VSV-EBO. Tissue samples (spleen, liver, kidney, and brain) were collected in medium-containing Precellys tubes, and blood was collected in tubes containing either EDTA, or without for serum collected at experimental endpoint which was 3 dpi for VSV-EBO infected animals, and 4 dpi for VSV-BDB infected animals.

#### Hyperimmune sera

*Ifnar*^*–/–*^ mice, (6–8-week-old) were intraperitoneally vaccinated on days 0, 7 and 14 with 5000 PFU of YF17D (n = 3), YF-EBO (n = 10), YF-SUD (n = 4), and YF-BDB (n = 5). Mice were bled via the submandibular vein on days 21 and 28, and were euthanized on day 35, after which blood was collected by cardiac heart puncture. Blood samples from individual mice were pooled together per group.

### Titrations

Tissue samples were homogenized in 350 µl of MEM assay medium using a Precellys 24 Touch (Bertin Technologies). The resulting homogenates were subjected to 10-fold serial dilutions, starting at a 1:10 ratio, and added in triplicate to 96-well plates pre-seeded with 2 × 10^4^ Vero E6 cells per well. Following a 6-day incubation in a humidified atmosphere (37 °C, 5% CO2), viral cytopathic effect (CPE) was quantified by staining viable cells with 0.2 mg/mL dimethylthiazol-carboxymethoxyfenyl-sulfofenyl-tetrazolium (MTS; Gibco) diluted in 1x MEM.

After a 1 h incubation, absorbance was measured at 498 nm using a SPARK Magellan plate reader (Tecan Life Technologies). The 50% tissue culture infectious dose (TCID50) for each sample was calculated according to the Reed and Muench method [45].

### Indirect immunofluorescent assay

HEK293Ad cells were seeded at a density of 2 × 10^4^ cells/well in 96-well plates 24 h prior to transfection. Cells were transfected with 0.25 µg/well of pCMV-GP-IRES-RFP plasmids expressing either SUDV, EBOV, BDBV, or RABV glycoproteins. At 24 h post-transfection, cells were fixed with 4% PFA, washed three times with PBS, and permeabilized with 0.1% Triton-X (Sigma) in PBS. Following a 30-minute blocking step (3% BSA in PBS) at room temperature, two-fold serial dilutions of mouse sera (starting at 1:40) were added and incubated for 1 h with orbital shaking (200 rpm). Bound antibodies were detected using a secondary goat anti-mouse Alexa Fluor 488 (AF488) antibody (Life Technologies) incubated for 1 h at 200 rpm. Nuclei were counterstained with DAPI (Sigma Aldrich, 200 µg/mL). Images were acquired using an Operetta CLS High-Content Imaging and Analysis system (Revvity, Inc.) equipped with a 10X objective. Appropriate filter sets were used for DAPI (Ex/Em: 350/470 nm), RFP (Ex/Em: 587/610 nm), and AF488 (Ex/Em: 493/518 nm). For each well, nine randomly selected fields were captured for quantitative analysis.

Image processing was performed using Harmony software (Revvity, Inc.). Viable cells were identified by DAPI-positive nuclei, with debris and non-viable cells excluded based on morphological and quantitative intensity thresholds. Transfected (antigen-positive) target cells were identified by the co-localization of DAPI and RFP signals. To determine endpoint titers while mitigating variability from non-specific serum binding, a texture-based analysis approach was employed. This method identified antibody-bound (AF488-positive) cells within the RFP-positive population based on morphological properties and signal distribution rather than absolute pixel intensity. This approach enabled the accurate and robust determination of binding antibody titers across different serum concentrations.

### Serum neutralization test

VeroE6-miRFP670– an in-house developed cell line expressing far red, using methodology described before [46] – were seeded in 96-well plates (2 × 10^4^ cells/well) and incubated overnight in a humidified incubator (37 °C, 5% CO_2_). Serial serum dilutions were prepared in a separate plate and incubated for 1 h with 7 PFU of VSV-BDB (replication-competent chimeric VSV) or at a concentration that yields around 10% transduction efficiency (SR-VSV), (37 °C, 5% CO_2_). Next, 25 µl of the virus-serum mixture was transferred to the cell-containing plates. After 32 h of incubation at 37°C with 5% CO_2_, the frequency of infected cells was quantified by imaging mCherry or GFP expressing cells using Harmony® software (version 5.2, LogiTech) on images acquired with Operetta CLS High Content Imaging system (Revvity). Half-maximal pseudovirus neutralization dilutions (PVNT_50_) were calculated following the protocol previously described [47].

### Statistical analysis and data representation

Statistical analyses were conducted using GraphPad Prism software (version 10.3.0, GraphPad Software, San Diego, CA, USA). For unpaired continuous variables, the Mann-Whitney U test was applied, while the Wilcoxon test was used for paired data. The Kruskal-Wallis test, followed by Bonferroni correction for multiple comparisons, was used to compare continuous variables across multiple groups. Survival data were evaluated using Gehan-Breslow-Wilcoxon test. Data are from distinct measurement and presented as mean ± standard error of the mean (SEM), unless otherwise stated. Differences with P < 0.05 (two-sided tests) were considered statistically significant. We did not assume normal distribution of datasets and therefore used non-parametric tests when comparing groups. Phylogenetic analyses and pairwise distances were constructed using MEGA12 software [48], using the Maximum Likelihood method based on the Tamura-Nei model for the phylogenetic tree and the Poisson model for pairwise distance.

## Data availability

All material and data supporting the findings in this study are available from the corresponding author upon reasonable request and considering third party rights under Material Transfer Agreement.

## Author contributions

Conceptualization: L.K., Y.A.A., K.D.; Investigation: L.K.; Methodology: L.K., V.L.; Formal analysis: L.K.; Validation: L.K., Y.A.A.; Visualization: L.K., Y.A.A.; Wrote the original draft: L.K.; Wrote the final draft: L.K., Y.A.A., K.D.; Supervised the study: J.N., K.D.; Funding acquisition: L.K., J.N., K.D.; Resources: J.N., K.D.; all authors approved the final manuscript.

## Acknowledgements

We are grateful to Prof. M. A. Whitt for sharing the original pVSV-ΔG-GFP-2.6 construct and helper plasmids, S. Debaveye for construction of expression plasmids, K. Geerts for expert help with *in vitro* virological and serological assays, W. Chiu and T. Francken for Vero E6-miRFP670 cells and help with analyzing high-content imaging data, C. De Keyzer for assistance with animal experimentation, K. van der Molen and F. Vanzeebroeck (Rega Institute Animal facility) for diligently taking care of animal breeding and husbandry. L.K. gratefully acknowledge funding support by a personal fellowship from the Flemish Research Foundation (FWO) (grant 1SH2H24N; NextEboVax). J.N. and K.D. acknowledge funding by the FWO Excellence of Science program (grant 40007527; VirEOS2) and the Rega Foundation, KU Leuven. K.D. acknowledges further support from KU Leuven Internal Funds (C24M/19/006), and funding by Horizon Europe, Health (grant agreement 101137459; Yellow4FLAVI), funded by the European Union. Views and opinions expressed are those of the authors only and do not necessarily reflect those of the European Union or European Health and Digital Executive Agency. Neither the European Union nor the granting authority can be held responsible for them. Part of this research work was performed using the ‘Caps-It’ research infrastructure (project ZW13-02) that was financially supported by the FWO Hercules Foundation and Rega Foundation, KU Leuven.

## Declaration of generative AI use

The authors report generative AI was not used in their research or preparation of this manuscript, with the exception of basic language assistance.

## EXTENDED DATA FIGURES

**Extended data figure 1.**
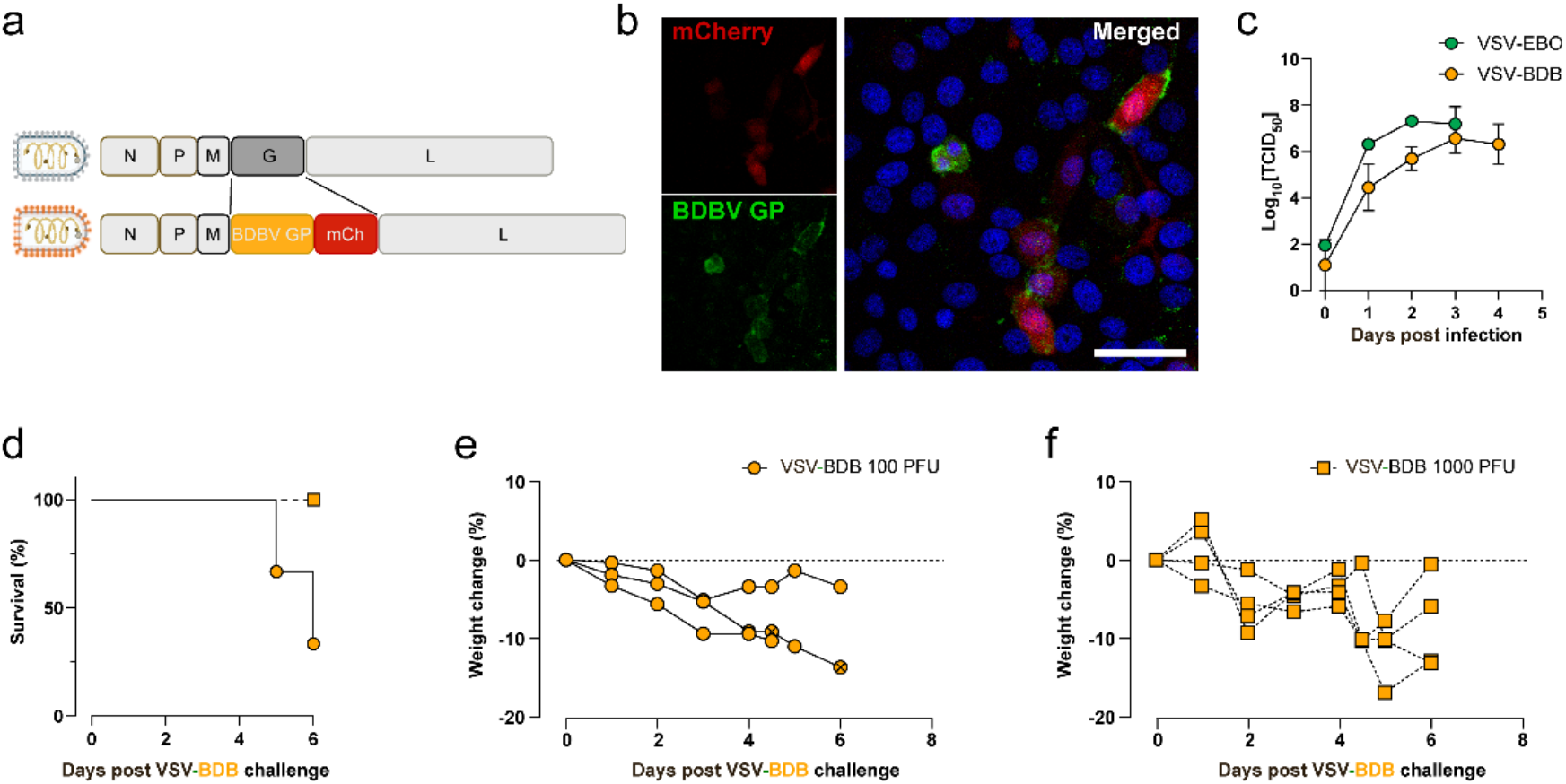
Characterization of the VSV-BDB infection model. **a**. Schematic representation of the recombinant VSV (rVSV) backbone, where the native VSV-G has been replaced with the Bundibugyo virus glycoprotein (BDBV GP). The construct includes an mCherry fluorescent protein reporter for the visualization of viral replication. **b**. Confocal imaging of Vero E6 cells infected with VSV-BDB, stained with polyclonal anti-YF-BDB serum (green), and nuclear stain (blue), while mCherry fluorescent protein from the virus is shown in red. Scale bar, 50 μm. **c**. *In vitro* replication kinetics of VSV-BDB compared to VSV-EBO in Vero cells (MOI = 0.01). Supernatants were harvested at indicated time points until total cytopathic effect (CPE) was observed. Data represent mean titers ± s.e.m. from two independent experiments. **d**. Survival (%) of naive *Ifnar*^−^/^−^ mice challenged intraperitoneally (i.p.) with either 100 PFU (n = 3) or 1000 PFU (n = 4) of VSV-BDB. **e, f**. Weight change (%) of naive *Ifnar*^−^/^−^ mice following infection with 100 PFU (**e**; n = 3) or 1000 PFU (**f**; n = 4) of VSV-BDB. Crossed datapoints indicate humane endpoints of individual animals.

**Extended data figure 2.**
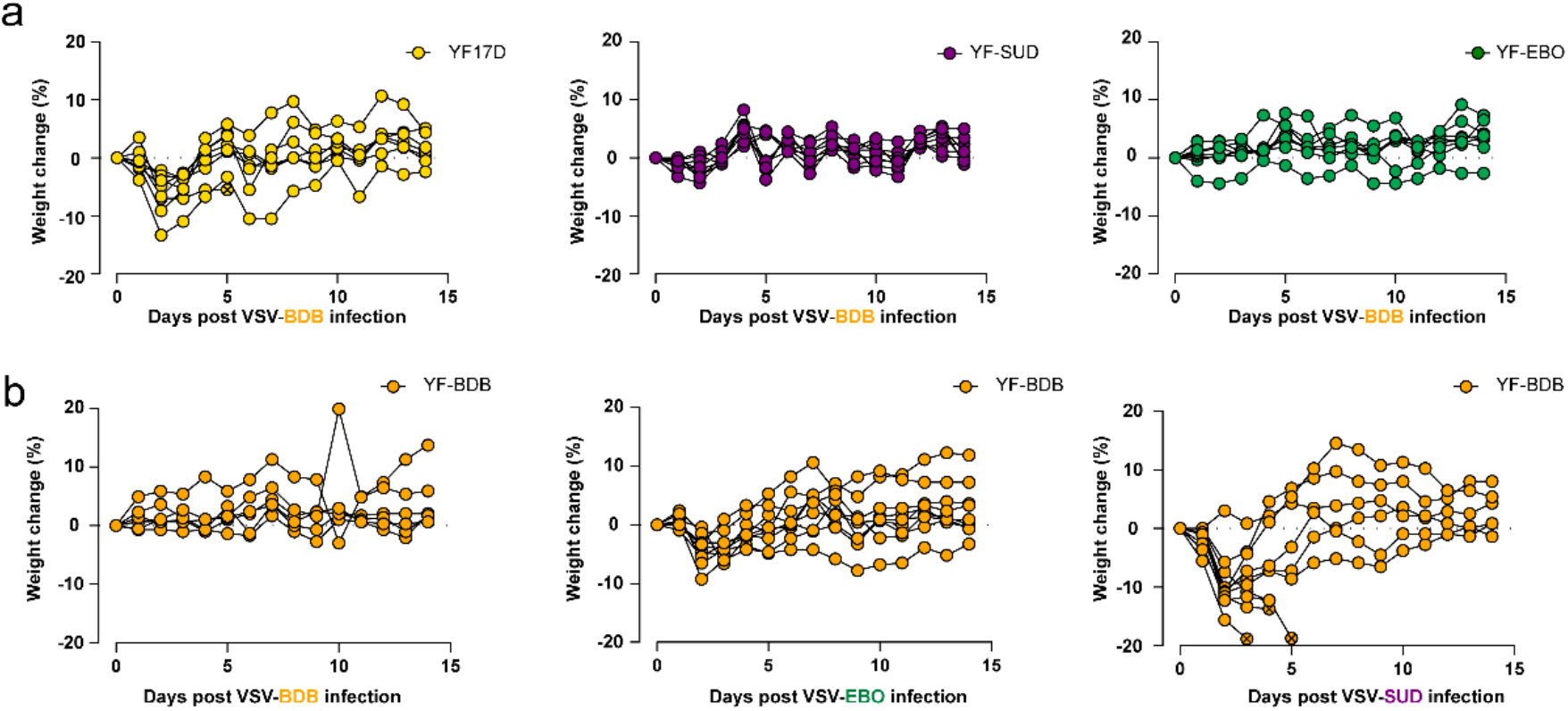
Weight change following VSV-BDB challenge and heterologous efficacy of YF-BDB. **a**. Weight change (%) of YF17D (n= 7), YF-SUD (n = 8), and YF-EBO (n = 7) vaccinated animals following challenge with VSV-BDB (100 PFU, i.p.). **b**. Weight change (%) of YF-BDB vaccinated *Ifnar*^−^/^−^ mice challenged with VSV-BDB (n= 6), VSV-EBO (n= 9), or VSV-SUD (n = 10). Crossed datapoints indicate humane endpoints of individual animals.

**Extended data figure 3.**
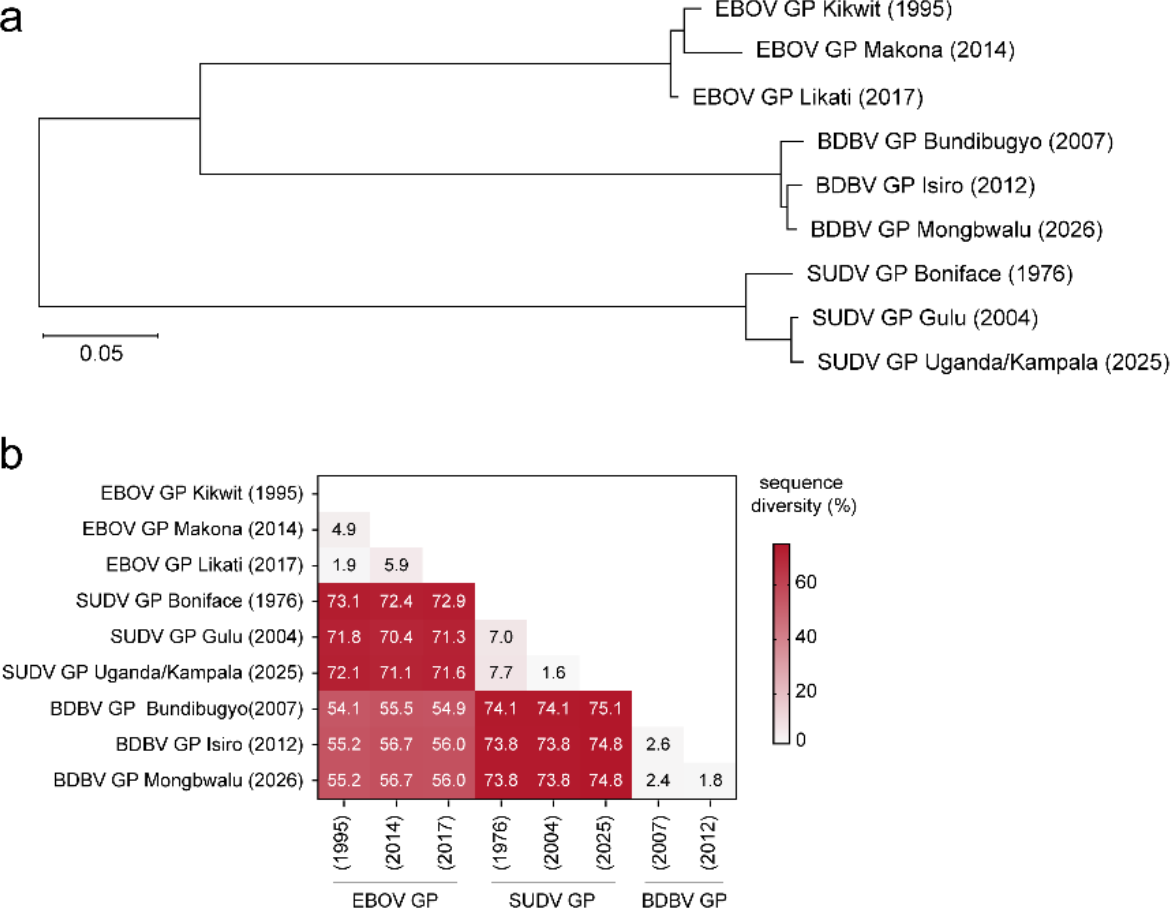
Phylogenetic relationship and GP sequence divergence among human-pathogenic orthoebolaviruses. **a**. Phylogenetic tree of orthoebolaviruses causing recurring outbreaks in humans, constructed using the Maximum Likelihood method based on the Tamura-Nei model. Branch lengths represent nucleotide substitutions per site. **b**. Pairwise amino acid distance matrix of the glycoprotein (GP) among representative *Orthoebolavirus* species. Divergence was calculated using the Poisson model. Phylogenetic analyses were done using MEGA12 software.

**Extended data figure 4.**
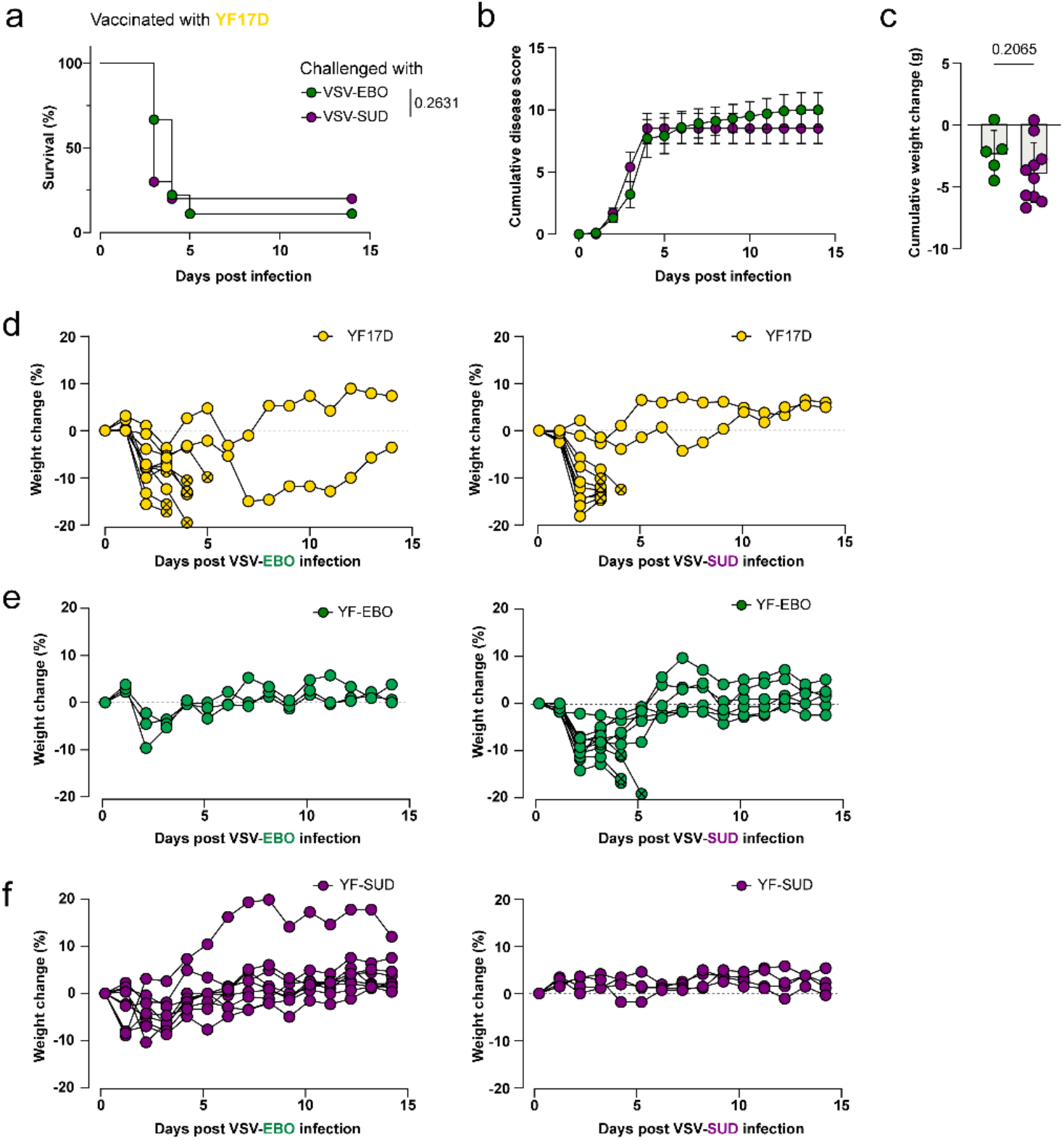
YF-SUD vaccination prevents severe morbidity following heterologous *Orthoebolavirus* challenge. **a-c.**Survival (%, **a**), cumulative disease score (**b**), and cumulative weight change (**c**) of YF17D-vaccinated mice following challenge with VSV-EBO (green, n = 10) or VSV-SUD (purple, n = 10). **d**. Weight change (%) of YF17D vaccinated (yellow, vector control) animals following challenge with either VSV-EBO (n = 10) or VSV-SUD (n = 10). **e**. Weight change (%) of YF-EBO (green) vaccinated animals following challenge with either VSV-EBO (n = 3) or VSV-SUD (n = 10). **f**. Weight change (%) of YF-SUD (purple) vaccinated animals following challenge with either VSV-EBO (n = 9) or VSV-SUD (n = 4). Crossed datapoints indicate humane endpoints of individual animals.

